# Advanced Bioprinting of Hydrogels with Controlled Mineral Gradients for Regenerative Engineering of the Osteochondral Interface

**DOI:** 10.1101/2024.11.25.625326

**Authors:** Xiao Zhao, Weiwei Wang, Xiaojun Yu, Dilhan M Kalyon, Cevat Erisken

## Abstract

The osteochondral (OC) interface exhibits a mineral gradient in the subchondral bone and articular cartilage interface, varying in thickness by several hundred micrometers across different species. Disruptions to this interface can cause severe damage to OC tissues, leading to osteoarthritis (OA), a debilitating and irreversible condition. Regenerative engineering approaches hold promise for addressing this issue by replicating the natural architecture and composition of native OC interface within a biomaterial scaffold. This study introduces a novel one-step bioprinting process using a twin-screw extruder that facilitates the fabrication of a unitary synthetic graft (USG), which mimics the native OC interface’s mineral concentration gradient.

The newly developed USG is composed of an agarose-based cartilage layer and a bone layer, which consists of agarose enriched with 20% hydroxyapatite (w/vol). The USG features a gradient interface with the mineral concentration seamlessly transitioning from 0 to 20wt% from the cartilage to the bone layer. The mineral gradients in the USG and the native tissue were documented using thermogravimetric analysis (TGA), micro-CT, and energy dispersive x-ray (EDX). TGA revealed that the gradient transition length in the graft (647±21μm) compared well to that of native OC tissue (633±124μm) harvested from bovine knee. The strain sweep and frequency sweep tests in oscillatory shear evaluated the linear viscoelastic properties of the grafts, indicating a dominant storage modulus over loss modulus similar to that of native OC tissues. Additionally, the compressive and stress relaxation behaviors of the USGs were quantified using multi-extensional tests, highlighting the grafts’ ability to maintain structural integrity under mechanical stress. Furthermore, viability assays performed after bioprinting showed that chondrocytes and human fetal osteoblast cells successfully integrated and survived within their designated regions of the graft. The USGs engineered in this study exhibit properties that make them promising candidates for regenerating OC defects and restoring knee joint functionality.

## 1. Introduction

Knee joint function is frequently compromised by a variety of causes including osteochondral (OC) tissue damage, which can arise from factors such as age-related degeneration, mechanical trauma, and other pathological conditions^1^. Untreated OC damages result in osteoarthritis (OA), a disease mostly associated with people over the age of 50, with an incidence of more than 250 million people worldwide^2^ and an economic burden of around 136.8 billion dollars^3^.

OA is a pathological process that cannot be reversed easily, mainly due to its avascular nature. This characteristic of the cartilage tissue significantly reduces its healing or regeneration capacity. Currently, the treatment approach for OA is a surgical procedure that includes but is not limited to microfracture, mosaicplasty, and ECM-based or synthetic grafts.^4^ Unfortunately, the outcome is generally a poor biological connection between the graft and the native tissue or the formation of fibrocartilage, which is weaker than its native counterpart both compositionally and functionally. Therefore, there is an unmet demand for designing and fabricating new synthetic or biological grafts that are similar to native tissue, i.e., biomimetic, in terms of composition, structure, and function.

The OC tissue is seen at the end of long bones as three different yet continuous layers of unmineralized cartilage (articular cartilage), mineralized cartilage (calcified cartilage), and subchondral bone^5,6^ or, more realistically, a graded structure from articular cartilage to subchondral bone^7,8^ (Figure 1).

**Figure 1.**
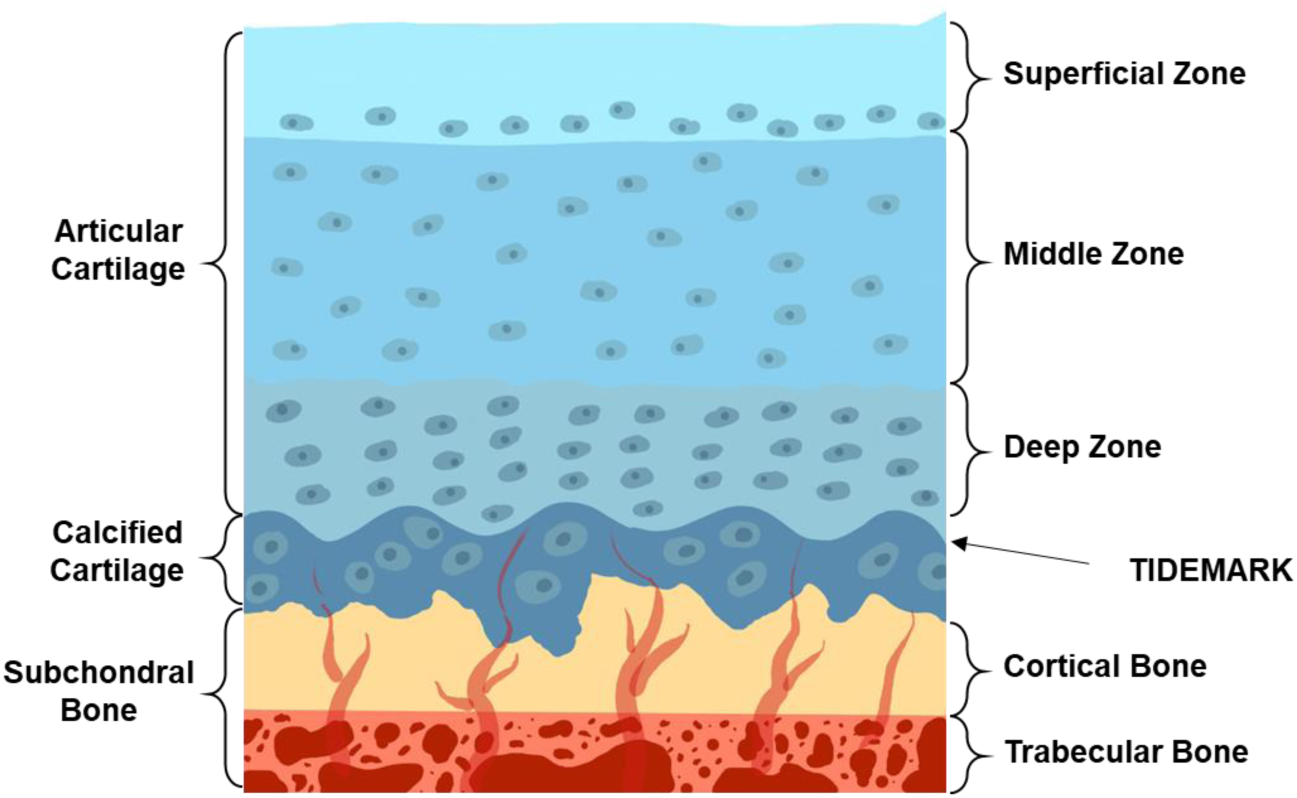
Structural representation of the osteochondral tissue (layer thicknesses are not true scale)

Successful replication of complex gradations such as those seen in the OC tissues requires the utilization of equally complex technologies to fabricate gradations in grafts. Such technologies can successfully address gradations seen in human tissues and tissue-tissue transitions, such as the interfaces between cartilage and bone, ligament and bone, and tendon and bone.^9–12^ Some of these technologies are also amenable to scaling up and fabrication at industrial levels under controlled and reproducible conditions. For example, a number of extrusion-based fabrication methods have been demonstrated for the fabrication of graded scaffolds in published reports.^13–16^

Various biocompatible materials, including natural polymers, synthetic polymers, and their combinations, have been employed to fabricate scaffolds for osteochondral tissue.^17^ Amongst these, agarose is a well-characterized hydrogel and has been widely used as a matrix for chondrocyte bioactivity and cartilage tissue engineering.^18,19^ Moreover, physiologically relevant mechanical properties that approach those of the native cartilage have been reported with chondrocyte-laden agarose constructs.^20^ Hydroxyapatite is present in bone tissue, and it is commonly used in bone tissue regeneration. Incorporating hydroxyapatite into hydrogels was shown to improve the mechanical properties and bioactivity of 3D printed scaffolds.^21,22^

Accordingly, the aims of this study were i) characterize the OC interface of the bovine knee, focusing on its unique structural and mineral components, ii) develop a continuously graded USG that closely replicates the structural properties of the native OC interface using agarose and hydroxyapatite as the primary biomaterials, and iii) employ extrusion bioprinting technology to fabricate the USG, ensuring it replicates the gradient of calcium content observed in natural OC tissues. We hypothesized that the hydrogel-based graft, created through advanced extrusion bioprinting techniques, will closely resemble the structural and compositional attributes of the native OC interface. This research is pioneering in its approach to simulate the physiological dimensions of calcium gradients within OC tissues.

The anticipated impact of the study is to provide valuable insights into the application of biomaterials for OC repair and regeneration, relevant in both preclinical and translational settings. The expected outcomes include significant advancements in orthopedic research, potentially influencing both technological and socio-economic spheres within the field. A major anticipated result is the development of grafts that reflect the physiological and structural characteristics of native tissues to help address the widespread need for effective OC regeneration solutions among the rapidly-aging global population.

## 2. Materials and methods

This research study involves three major experimental studies: (i) harvesting and characterization of OC tissues from bovine knee joints, (ii) printing grafts and their characterization, (iii) comparing the two groups in terms of distribution of mineral content as well as biomechanical and rheological properties, and (iv) bioprinting chondrocytes and hFOBs in hydrogel and investigating their viability. The overall experimental procedure is shown in Figure 2.

**Figure 2.**
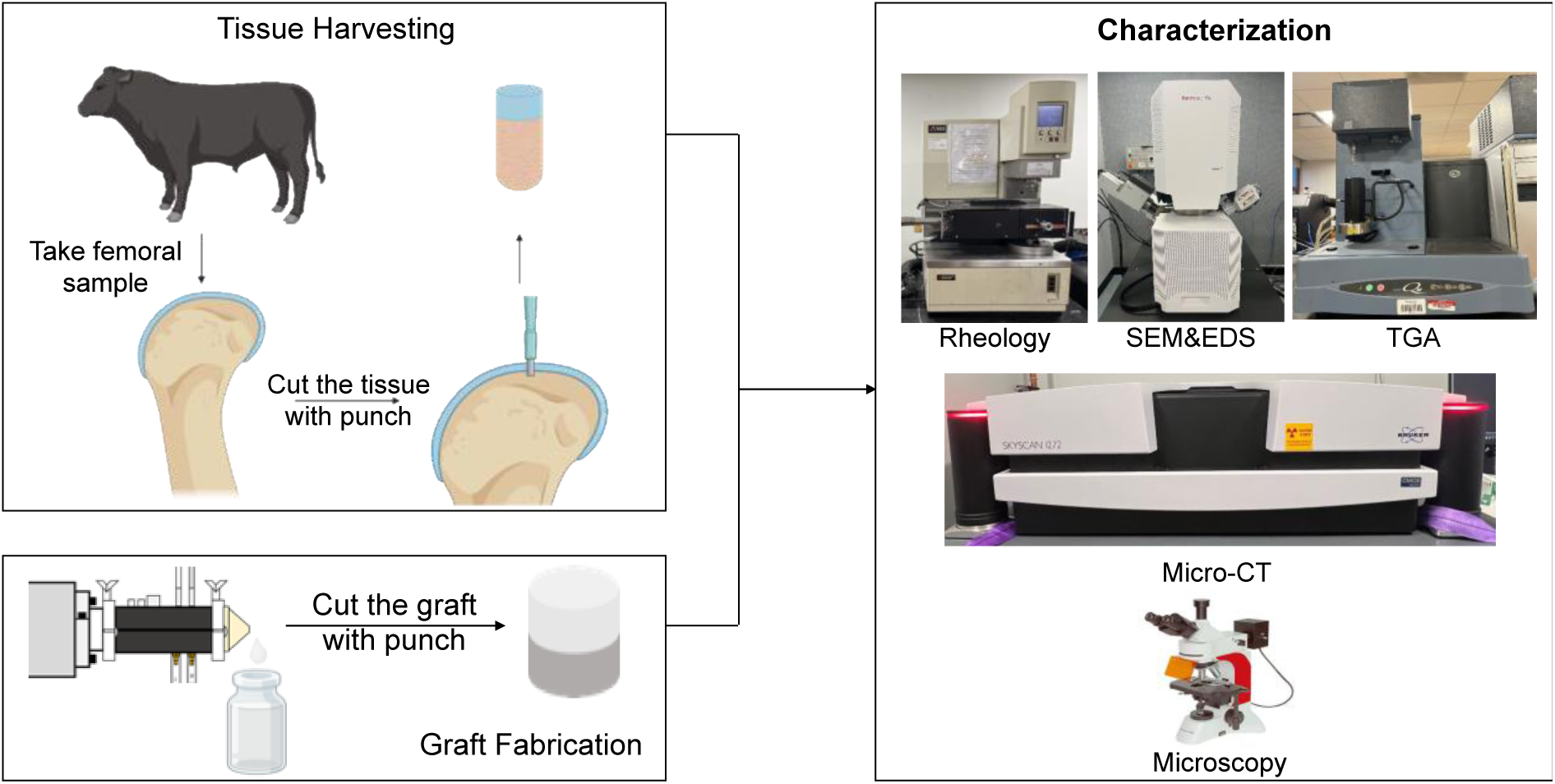
Experimental procedures for the tissue harvest, graft fabrication, and characterization.

### 2.1 Materials

A low-melting-temperature agarose was obtained from Cambrex Bio Science Rockland (Product #50100, Rockland, ME). Hydroxyapatite (Product #04238) and phosphate-buffered saline (PBS, Product #524650) were sourced from Sigma-Aldrich, both located in Saint Louis, MO. Detailed sources of biological materials are specified in the appropriate sections of the manuscript.

### 2.2 Harvesting the OC Tissue

The bovine knee joints (1-2 year old, n=3) were obtained from a local slaughterhouse and temporarily preserved at 4°C until needed. Before testing, the refrigerated knee joints were brought to room temperature, and plugs were harvested from the femoral condyle using punches with diameters of 4mm and 10mm (Figure 3). Specimens with a 4mm diameter were utilized to evaluate mineral content using a thermogravimetric analyzer (TGA), while those with a 10mm diameter were used for micro-CT, biomechanical, and rheological measurements.

**Figure 3.**
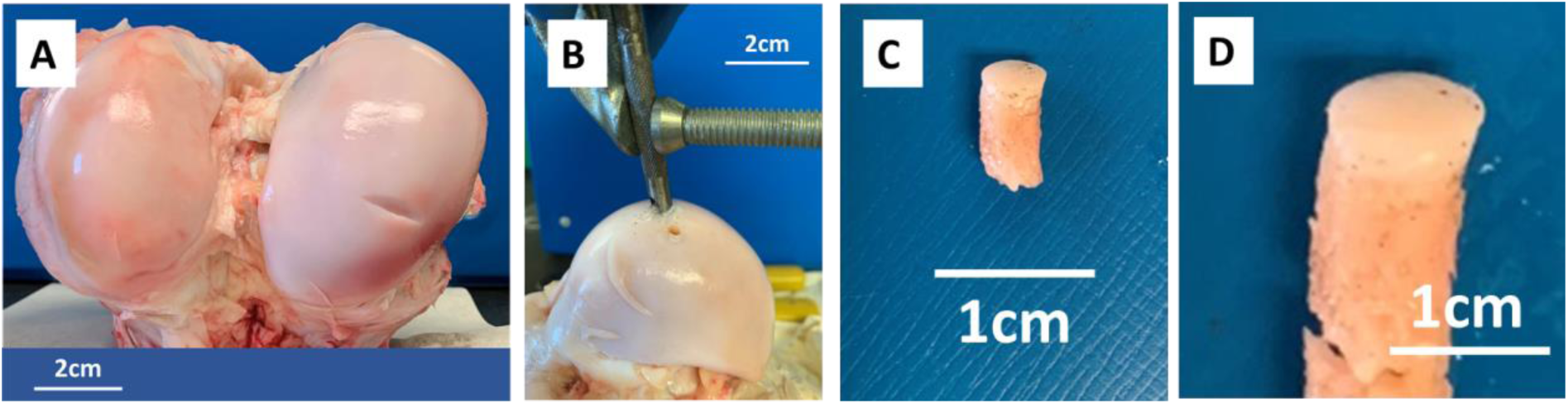
Harvesting and preparation of OC tissue for characterization. (A) Gross view of femoral condyles harvested from bovine, (B) specimen collection using 4mm and 8mm diameter punches, and specimens for (C) thermogravimetric and (D) micro-CT, biomechanical, and rheological characterization.

### 2.3 Formulation and extrusion printing of graded OC grafts

The USG was engineered to incorporate a chondrogenic region, a transition region, and a bone region, and was fabricated accordingly. The chondrogenic hydrogel was prepared by dissolving agarose in phosphate-buffered saline (PBS) to achieve a 2% concentration (0.02 g/mL). The solution was heated to boiling three times to ensure complete polymerization. The osteogenic hydrogel was formulated by adding hydroxyapatite nanoparticles to the 2% agarose solution at a 20% (0.2g hydroxyapatite per mL of 2% agarose) concentration. The hydroxyapatite was sonicated post-mixing to disrupt any agglomerates that may have formed. Both the agarose and agarose/hydroxyapatite mixtures were non-flowable at room temperature but transformed into injectable fluids at and above 37.5°C, enabling their administration into the extruder via infusion pumps. The osteochondral USGs were finalized by cooling to room temperature, where both the agarose solution and agarose/hydroxyapatite suspension solidified into gels (see Figure 4).

**Figure 4.**
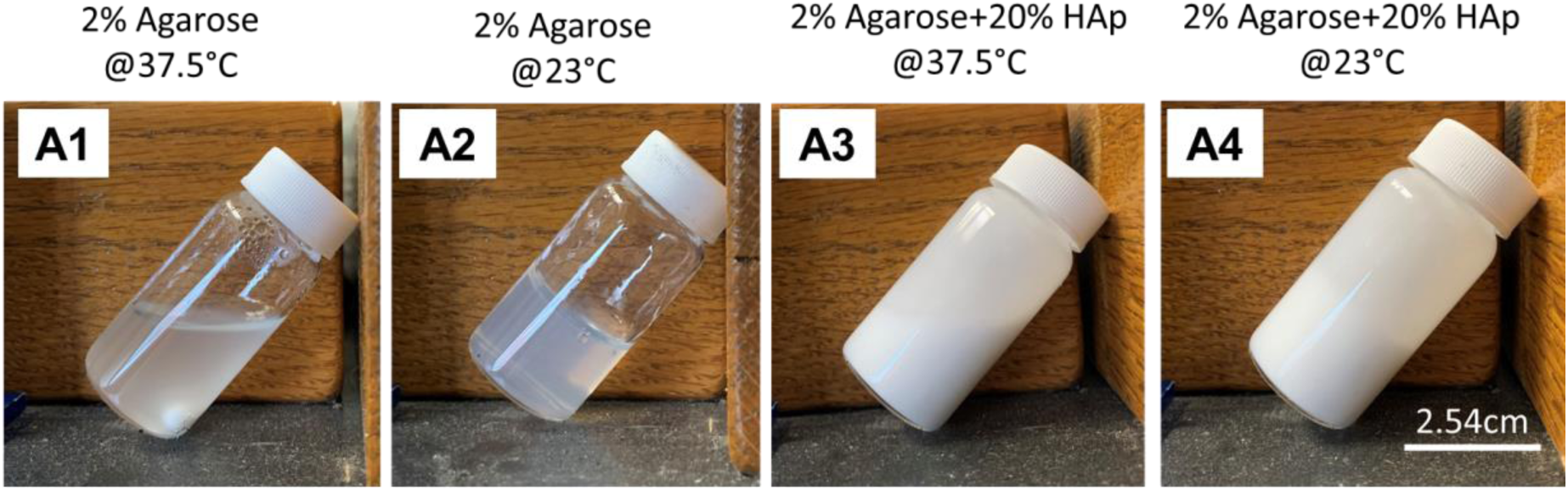
Formulation of hydrogel for synthetic graft fabrication. The 2% agarose was formulated to be injectable at 37.5°C (A1), and to exhibit gel-like behavior at room temperature (A2). Similarly, the suspension prepared with 20% (by weight) hydroxyapatite, HAp, was injectable at 37.5°C (A3), and gel-like at room temperature (A4).

The fabrication of functionally graded biomimetic USGs primarily involves a twin-screw extruder with fully intermeshing, co-rotating screws of 7.5 mm diameter, custom-built by Material Processing and Research, Inc., of New Jersey. This extruder is equipped with a slit die, tailored specifically for this application (refer to Table 1 for die dimensions). Multiple feeding ports enable the continuous introduction of various solutions/suspensions, such as the agarose solution and agarose/hydroxyapatite suspension used in this study. The screws of the extruder are designed to enhance dispersive mixing, effectively capable of breaking up any agglomerates of hydroxyapatite that form during initial mixing. This capability was further customized by employing combinations of kneading discs and fully flighted elements,^23–26^ which are staggered at specific angles and directions to optimize distributive and dispersive mixing efficiency.

**Table 1:** Parameters used to optimize the gradient transition length.

| Run # | Lag time (s) | Rotational speed (RPM) | Die opening (mm x mm) (Width x Height) |
| --- | --- | --- | --- |
| 1 | 30 | 50 | 10x3 |
| 2 | 12 | 50 | 10x3 |
| 3 | 0 | 75 | 2x2 |

The twin screw extruder was housed within an environmental chamber maintained at 37.5°C, while the barrel temperature was controlled at 37.5°C by circulating water through a heat bath (see Figure 5 A, B). This setup prevented the premature gelation of the agarose and agarose/hydroxyapatite suspensions during processing. The agarose solution was introduced through an injection point near the die, and the agarose/hydroxyapatite suspension was fed through another injection point. The capability to adjust the feed rates of agarose and agarose/hydroxyapatite in a time-dependent manner enabled the creation of a continuously varying hydroxyapatite concentration within the osteochondral USG, ranging from 0–20% by weight. The overall feed rate to the extruder was maintained at 10 mL/h. The feed rates for agarose and agarose/hydroxyapatite were incrementally adjusted by 1.5 mL/h intervals, with one increasing while the other decreased to maintain a constant total rate, thereby establishing a graded structure. Specifically, the flow rate of agarose was incrementally increased from 0 mL/h to 10 mL/h, while that of agarose/hydroxyapatite was decreased from 10 mL/h to 0 mL/h. Additionally, the gradient formation within a specific length was controlled by varying the lag time between feed rate adjustments, the rotational speed of the screws, and the die opening. The impact of these adjustments on the gradient distance is detailed in Table 1, which lists the optimized parameters for achieving the desired grading distance.

**Figure 5.**
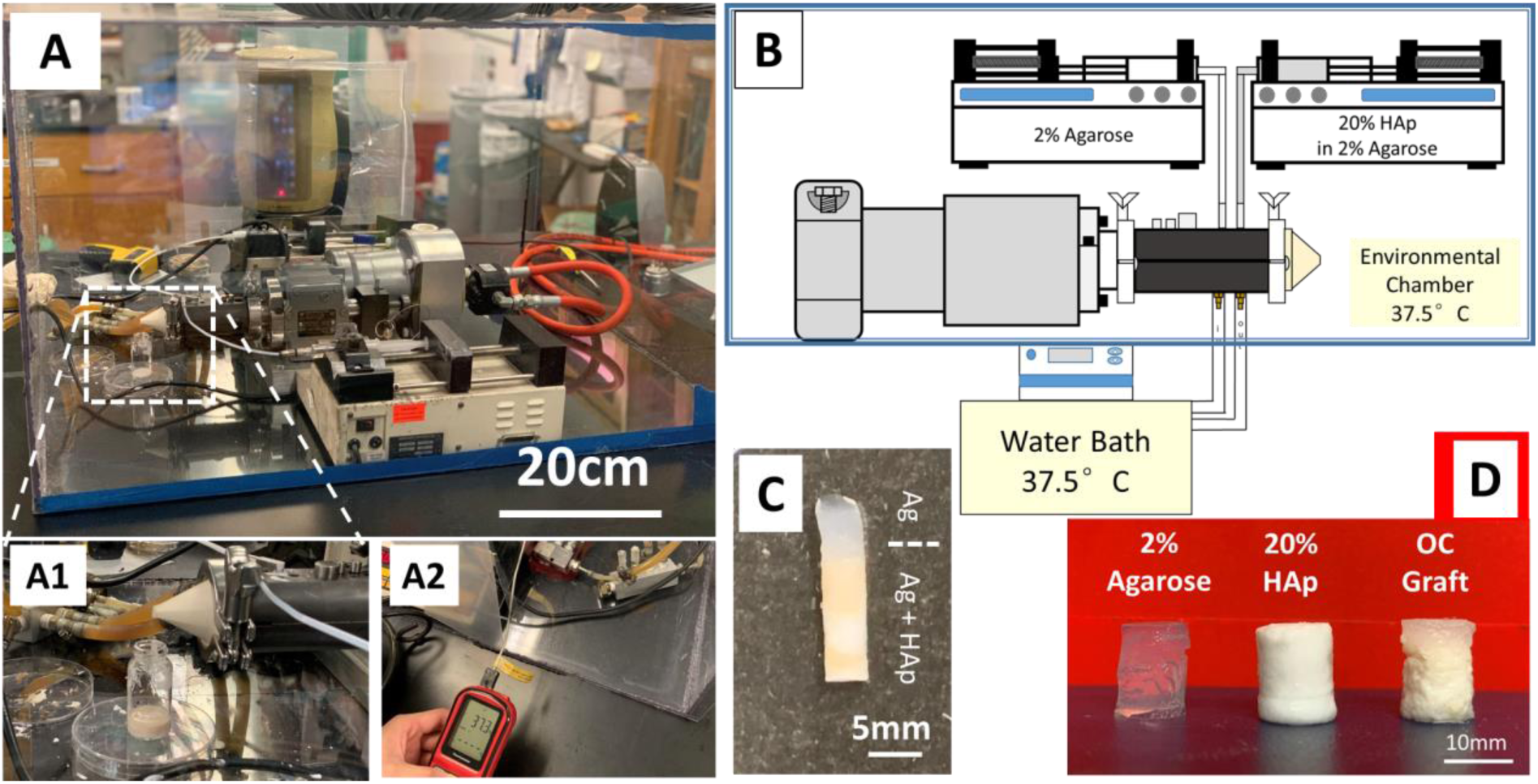
Extrusion printing of OC grafts. OC grafts were fabricated to contain hydroxyapatite at the bottom part of the grafts, a continuously decreasing concentration from 20% to no-hydroxyapatite, and agarose-only at the top to mimic bone, OC interface and cartilage, respectively. Grafts were fabricated using a twin-screw mini extruder installed in an environmental chamber (A, B), where 2% agarose solution and agarose/hydroxyapatite were fed from two separate infusion pumps with changing flow rates to create gradients of hydroxyapatite. The grafts fabricated were, then, punched to have diameters of 4mm (C) and 8mm (D) for thermogravimetric, rheological, and micro-CT characterizations. HAp: hydroxyapatite.

The combined agarose and agarose/hydroxyapatite with the conditions defined in Table 1 were extruded into vials 25mm in diameter, and the vials were moved from the environmental chamber to room temperature to eventually form USG. Finally, these grafts were punched into cylindrical shapes for thermogravimetric, mechanical, rheological, micro-CT characterization, and microscopic visualization. For SEM and EDX characterizations, the specimens were cut into rectangular shapes instead of cylindrical shapes for better imaging.

### 2.4 Thermogravimetric characterization of OC tissues and grafts

The weight fraction of mineral content in the native OC tissues and the hydroxyapatite content in the OC USGs were validated by using a thermo-gravimetric analysis apparatus (TGA-Q50, TA Instruments of New Castle, Delaware). The tissues harvested from femoral condyles and the USGs were stored in sealed packages at 4°C until needed and kept in a hydrated chamber during sample collection to prevent dehydration. Slices were cut from the cylindrical tissues and grafts, their thickness was measured using a digital caliper (AOS 500, Mitutoyo, Japan), and the sections were placed in the analyzer’s sampler. The residue of the sliced specimen was then determined upon heating from 25 to 550 °C at a rate of 25 °C/min under nitrogen.

### 2.5 Scanning Electron Microscope and EDX characterization

The OC tissues and USGs were freeze dried overnight, coated with gold and imaged using a Thermo Fisher Scientific Apreo 2 scanning electron microscope (SEM). Energy dispersion X-ray (EDX) and energy dispersive spectroscopy (EDS) of the OC grafts were also conducted under SEM upon scanning for 300s duration at 20 kV to validate the presence of calcium and phosphorus.

### 2.6 Biomechanical characterization of OC tissue and grafts

The compressive properties of OC tissues and OC USGs were characterized using a Rheometric Scientific ARES Rheometer (currently TA Instruments) in the strain range of 0%– 5% at a constant compression rate of 0.05 mm/s while immersed in a phosphate-buffered saline (PBS) environment at 37°C. This strain level was previously used in biomechanical characterizations of native cartilage and engineered tissues.^7,27^ Here, the specimens were initially squeezed using a normal force of approximately 0.03N to obtain full contact, and the compression test was then initiated.

### 2.7 Linear viscoelastic properties and relaxation of OC tissue and grafts

The samples were characterized in compression and oscillatory shear using the ARES rheometer (TA Instruments, New Castle, DE). Briefly, the specimen was inserted between 2 disks (8-mm diameter), which were also immersed in PBS solution kept at 37°C to prevent drying of the specimens of the native tissue and the grafts during the experiments^27^. The upper disk either oscillates in the clockwise and counterclockwise directions or translates in the downward direction at a constant velocity, whereas the second disk remains stationary and is connected to the torque and normal force transducer.

In small-amplitude oscillatory shear, the shear strain oscillates as a function of time as: γ=γ_0_ sin(ωt), where γ_0_ is shear strain amplitude (i.e., θD/h, where θ is the angular displacement, D is the disk diameter and h is the gap between the two disks), ω is the oscillation frequency, and *t* is time. The shear stress, τ, response to the oscillatory deformation consists of two components related to the energy stored and energy dissipated as heat, i.e., τ =G′(ω) γ_0_ sin(ωt)+G″ (ω) γ_0_ cos(ωt), where G′ (ω) is the shear storage modulus and G″ (ω) is the shear loss modulus. The ratio of G″ (ω) to G′ (ω) is tanδ. The oscillatory shearing needs to be carried out in the linear viscoelastic region at which the moduli are independent of the strain amplitude. The strain amplitude sweeps indicated that up to a strain amplitude of 0.1%, the oscillatory shear deformation of the hydrogels and the native tissue occurred in the linear region. The dynamic properties G′ (ω) and G″ (ω) were characterized as a function of frequency in the range of 0.1– 100 rps at 0.1% strain. Sweeping the frequencies enables the characterization of the linear viscoelastic response of the tissue over a range of time scales. In general, at relatively short characteristic times of deformation, the elastic response is emphasized, while the viscous flow behavior is emphasized at longer characteristic times. Time sweep tests performed earlier on cartilage tissue^27^ and hydrogels^28^ suggested that the samples were stable within the time scale of the experiments (typically less than 15 minutes of shearing for each sample).

### 2.8 Micro-CT characterization of OC tissue and graft**s**

The bovine osteochondral tissue samples were harvested from the bovine knee joint and tested immediately. The samples were kept hydrated with a customized moisture chamber during the testing. The graft specimens were tested after freeze drying in the air. The Micro-CT machine (SkyScan 1272, Bruker, MA, USA) was operated with a source voltage of 60kV, a source current of 70μA, 2×2 binning, Al filter of 0.5mm and a resolution of 2048×2048 with varied pixel sizes. The 3D reconstruction of bone mineral density was done with the built-in micro-CT software set from Bruker (CTVox and CTAn).

### 2.9 Bioprinting of human fetal osteoblasts (hFOB) and chondrocytes

#### Cell culture, growth and labelling

Human fetal osteoblast (hFOB, ATCC, Catalog#CRL-3602) cells and rat articular cartilage chondrocytes (Cells-Online LLC, CA, USA) were used for the bone and cartilage regions of the OC USG, respectively. The hFOB cell line was maintained in a 1:1 mixture of Ham’s F12 Medium and Dulbecco’s Modified Eagle’s Medium, supplemented with 2.5 mM L-glutamine, 1% antibiotics (penicillin/streptomycin), and 10% fetal bovine serum (FBS). The cells were cultured at 37 °C in a 5% CO_2_ incubator, and the medium was changed every 2 days. A total of 600K hFOB cells were labelled using Green CMFDA (Thermofisher, #C7025) and mixed with 20mL of agarose/hydroxyapatite suspension, and fed to the extruder unit for bioprinting. Rat articular cartilage chondrocytes were cultured in DMEM supplemented with 10% fetal bovine serum, and 1% penicillin/streptomycin. Cells were incubated at 37 °C and 5% CO_2_ in the incubator. Culture media was changed every other day. Agarose hydrogel scaffolds were prepared in sterilized PBS. A total of 300K chondrocytes were mixed with 10mL of 2% agarose (0.02g agarose / mL PBS), and fed to the extruder for bioprinting. Before infusing the hydrogels to the extruder, the chondrocytes were tested for viability in agarose of different concentrations (0.5-2%) for cell viability.

Cell tracker CMFDA (Thermofisher, #C7025) was allowed to warm to room temperature before breaking the seal. Then, Green CMFDA was dissolved in DMSO to a final concentration of 10 mM as a stock solution. The stock solution was diluted to a final working concentration of 5 µM for labeling. For labeling, culture media was aspirated, the pre-warmed working solution was added to the flasks, and the content was incubated for 30 min at 37 °C in a 5% CO_2_ incubator. Upon culture, the working solution was aspirated, and cell culture media was added as described above. The resulting cells were visualized through a fluorescent microscope (emission: 514 nm and excitation: 485 nm) for 72h.

After being embedded into the agarose gel, cells were bioprinted immediately.

#### Extrusion bioprinting of cells

The 2% agarose with chondrocytes and the 20% hydroxyapatite/agarose (0.2g hydroxyapatite per mL of 2% agarose) with hFOBs were fed to the extruder, and the cellular OC graft was formed under conditions described in Section 2.3. The bottom part of the bioprinted construct served as the bone part and contained hFOBs, and the top part was for cartilage and contained chondrocytes. Cylindrical specimens were punched and processed for staining and imaging.

#### Imaging cells and matrix in the OC graft

There is no additional staining of hFOB cells other than labeling. One drop of DAPI solution was added to the sections to stain the cell nuclei just prior to imaging.

For sectioning the OC grafts, the samples were incubated either in 15% (v/v) or 30% (v/v) sucrose in PBS and embedded in the optimal cutting temperature (O.C.T.) compound. The specimens were sectioned into 8-μm-thick slices with the cryostat microtome.

The mineralized matrix within the grafts was analyzed using Alizarin Red staining. Sections were rinsed with PBS, fixed in 10% neutral buffered formalin for 30 minutes, and stained with 2 mL of 40 mM Alizarin Red S (ARS) (Millipore, TMS-008-C) for 30 minutes under gentle agitation. The sections were then rinsed ten times with deionized water to remove non-specific binding. Imaging was conducted using a Nikon T100 Stereomicroscope.

Fluorescent imaging utilized CMFDA for green staining and DAPI for blue staining of the cell nuclei. This was performed using a Nikon SMZ1500 stereomicroscope.

### 2.10 Statistical analysis

The thicknesses of the gradients found in the native OC and the grafts were compared using the Student’s-t test. The modulus of the graft and the first and second moduli of the native tissue (which exhibited two distinct moduli) were compared using one-way ANOVA. Statistical significance was established at p < 0.05.

## 3. Results

### 3.1 Gradient transition length as determined by TGA

The gradient transition length describes the length over which the hydroxyapatite concentration changes from high to low (from bone to cartilage). Two sets of experiments were performed using thermogravimetric analysis. First, the mineral content of the bovine OC tissue was analyzed across the bone-cartilage interface. In a similar set of experiments, the variation in hydroxyapatite concentration across the bone to cartilage transition in the printed USGs was quantified. The harvested bovine OC tissues were around 10 mm in overall thickness. Given that the cartilage and OC layer together measure approximately 2 mm, assessments were conducted up to 3 mm from the articular surface.

Based on TGA measurements, the gradient transition length for the native OC tissue was found to be 633±124 µm (Fig 6 A, D, n=3). The printing of the USGs under conditions specified in Table 1 generated grafts with different gradient transition lengths (Fig 6B, D). Based on adjustments to the flow rate interval, reducing the time between flow rate changes from 30 seconds to 12 seconds decreased the gradient transition length from 3830 µm to 2720 µm. Instantaneously altering the flow rate, alongside increasing the screw rotational speed and reducing the die opening, further decreased the gradient transition length to 647 ± 21 µm. As shown in Figure 6C, the gradient transition lengths of the OC tissue and the USG, measured using TGA characterization, were comparable (p > 0.05).

**Figure 6.**
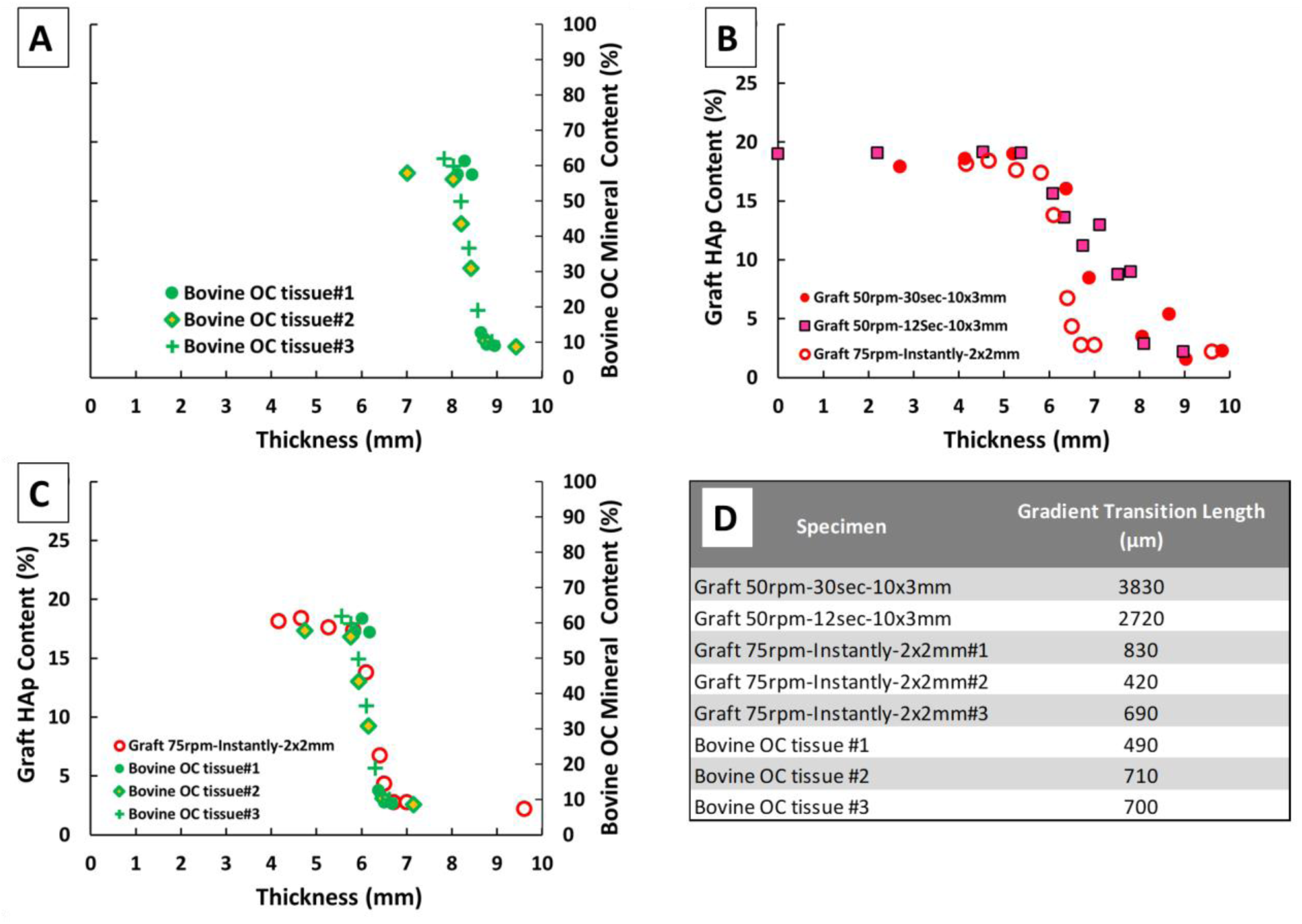
Determination of mineral gradient in bovine OC tissue (A), grafts (B) and comparison of the two (C, D). (A) The mineral concentration of the bovine OC tissue (∼10mm thick) was determined as a function of the thickness of the tissue (n=3). The bottom and top part of the OC tissue corresponds to bone and cartilage tissues, respectively. (B) Process parameters (screw rotational speed and lag time) and extruder design parameters (die dimensions) were changed to form a gradient of hydroxyapatite (HAp) concentration in the graft similar to that seen in native OC tissue. (C) The gradient transition lengths in OC tissue and the graft were overlapped. (D) The gradient transition lengths for grafts fabricated under different conditions and for the bovine OC tissue.

### 3.2 SEM and EDX characterization

The SEM imaging, EDX elemental mapping, and line scan techniques demonstrated the presence of grading in both native OC tissue and the USGs of this study qualitatively and semi-quantitatively. The change in tissue and graft composition as a function of distance from the surface was apparent in the form of color difference in gross images (Fig 7A1, B1). Structural and compositional (as calcium content) changes were also demonstrated in SEM (A2, B2) and EDX (A3,4 and B3,4) characterizations.

The results showed that calcium was absent on the surfaces of the OC tissue and the USG, and the concentration of calcium gradually increased in the region of cartilage-bone transition, after which it remained constant. This change in calcium content was quantified and shown in Fig 7C1. Clearly, both OC native tissue and graft exhibited a gradual change in calcium content at the interface region.

**Figure 7.**
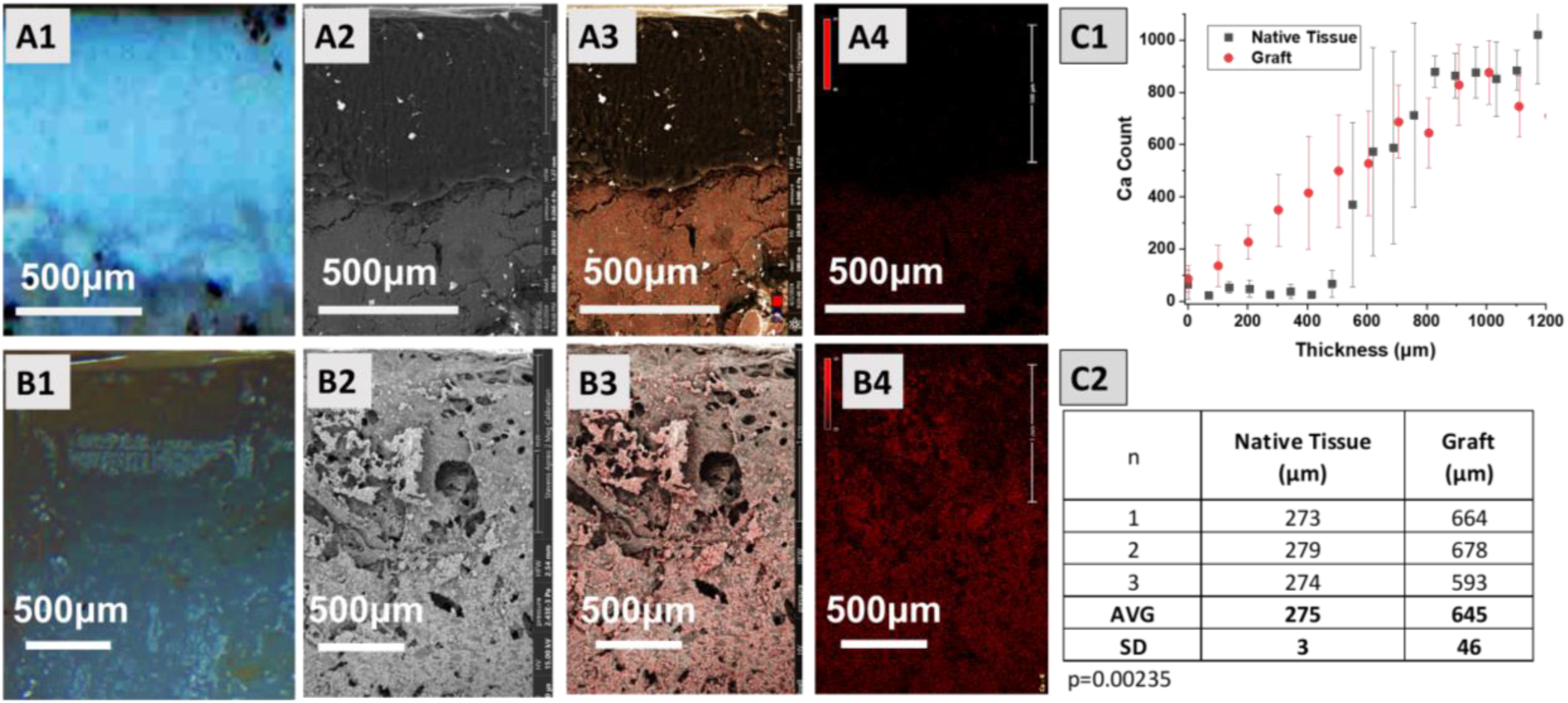
Gross image (A1), SEM micrograph (A2), and calcium mapping (A3-A4) of fresh bovine OC tissue compared with gross image (B1), SEM micrograph (B2), and calcium mapping (B3-B4) of the OC graft (USG) of this study. (C1) The grading distance for native OC tissue and grafts as determined by line scans over the EDX mapped images and (C2) corresponding gradient lengths (n=3). Error bars represent SD.

### 3.3 Linear viscoelasticity and biomechanical properties of OC tissue and grafts

The linear viscoelastic material functions of both the USGs and the native OC tissues were characterized using an ARES rheometer. Oscillatory shear experiments were conducted to evaluate sample stability under shear, determine the strain amplitude range for linear behavior, and explore the dependence of storage and loss moduli on deformation rates (frequency).

Initially, a time sweep test lasting approximately 15 minutes was conducted to verify data reliability. The results indicated stable properties, with only the storage modulus reported, over a 900-second duration (Figure 8A). We also assessed the temperature sensitivity of the tissues and grafts between 19°C and 39°C, observing no significant effects (Figure 8B).

**Figure 8.**
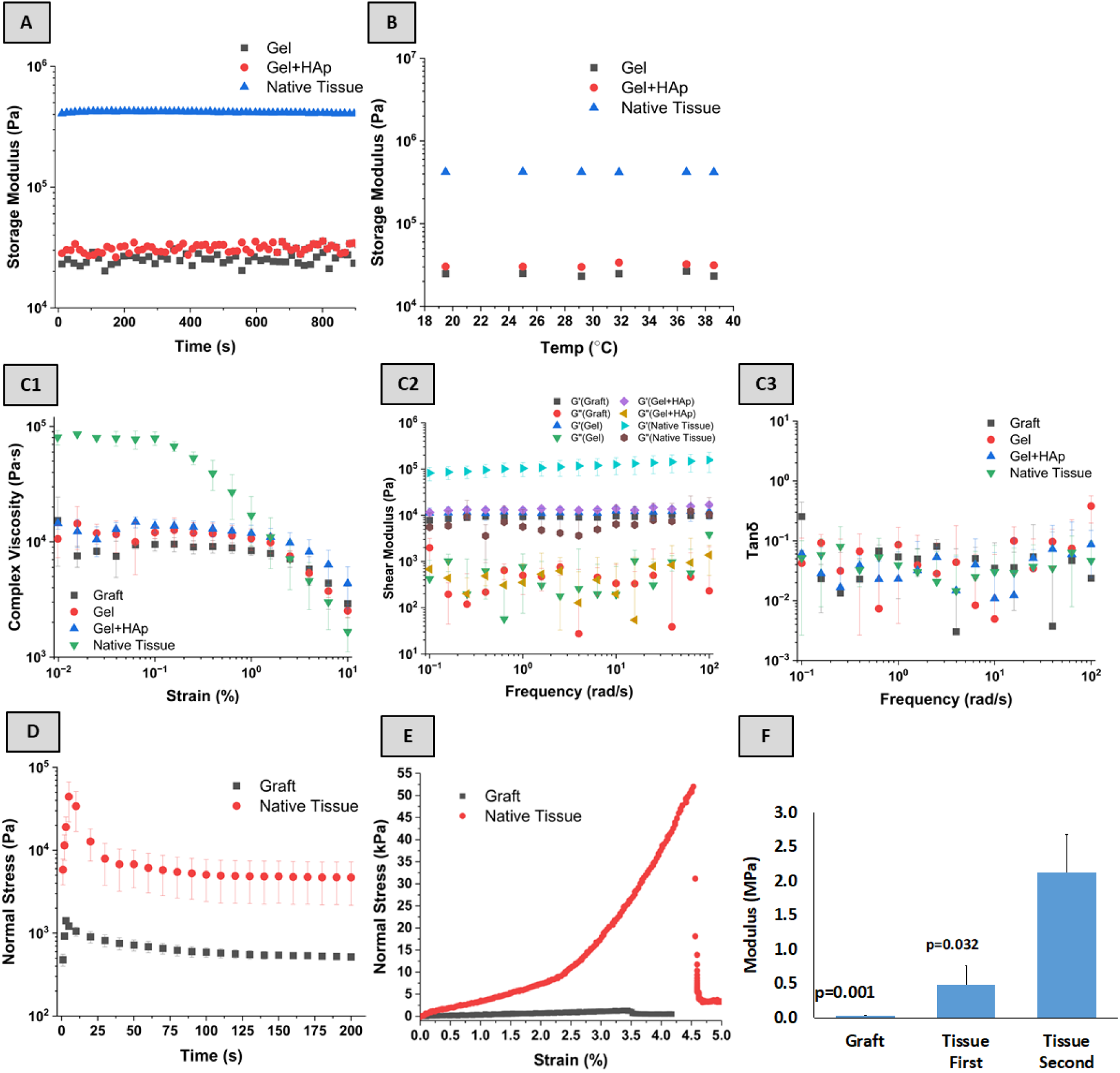
Linear viscoelastic material functions and compressive properties of bovine OC tissue and USG (graft). (A) Time sweep of the native tissue, agarose (gel) and agarose+hydroxyapatite (Gel+HAp); (B) temperature sweep of the OC tissue, agarose and agarose+hydroxyapatite; (C1) strain sweep (at 1rps), (C2) frequency sweep (at 0.1%) behavior of native OC tissue, agarose, agarose+hydroxyapatite and USGs at 37°C, (C3) tan δ; (D) relaxation behavior of the OC tissue and USGs; (E) Stress-strain behavior of the USG and OC tissue with two moduli; (F) compressive moduli of the OC tissue and graft. Error bars represent standard deviation. The p values show a comparison with the second modulus of the OC tissue.

Subsequently, a strain sweep test identified the linear viscoelastic range for each sample. The native OC tissues exhibited consistent rheological properties up to a 0.1% strain amplitude, whereas the grafts and their components (agarose and agarose/hydroxyapatite) maintained stable properties up to a 1% strain amplitude (Figure 8C1). This led to a frequency sweep test conducted at 0.1% strain, where the OC tissues and all graft compositions displayed typical gel-like behavior. This is indicated by frequency-independent responses in storage (G′) and loss (G") moduli (Figure 8C2, 3), with the storage modulus significantly exceeding the loss modulus, as reflected in tan δ values below 1 (Figure 8C3).

The compression stress relaxation behavior was analyzed by imposing a 5% compression over 2 seconds (see Figure 8D). Following the initial strain, the normal stress in both native OC tissues and grafts gradually decreased, exhibiting a time-dependent relaxation characterized by a gradual decrease in normal stress, which stabilized around 200 seconds. This similarity in behavior under compression between the native tissues and the USGs highlights the successful replication of the native tissue’s viscoelastic properties in the USGs.

The compressive stress-strain behavior of both the native OC tissue and the graft was rigorously evaluated. As depicted in Figure 8E, the native OC tissue exhibited dual moduli. The initial modulus, representing the cartilage region, was recorded at 0.48±0.28 MPa. The second modulus, indicative of the bone region, was significantly higher at 2.13±0.55 MPa (p<0.05 when compared to the initial modulus). Conversely, the OC graft displayed a consistent linear stress-strain behavior until failure. The modulus of the graft was notably lower, measured at 0.04±0.01 kPa (p<0.05 when compared to the second modulus of the native tissue), as illustrated in Figure 8F.

### 3.4 Micro-CT characterization of OC tissue and graft

Both native OC tissue specimens and grafts were examined using micro-CT imaging, and the mineral density was measured by dividing the entire specimen into layers with 12-14µm thickness. The images of the specimens clearly show the cartilage, bone, and the transition between the two (Fig 9 A, B). The change in bone mineral density of the specimens was plotted as a function of length from the surface as shown in Figure 9C1 for individual runs and in Figure 9 C2 in the form of means and deviations from the mean (SD).

**Figure 9.**
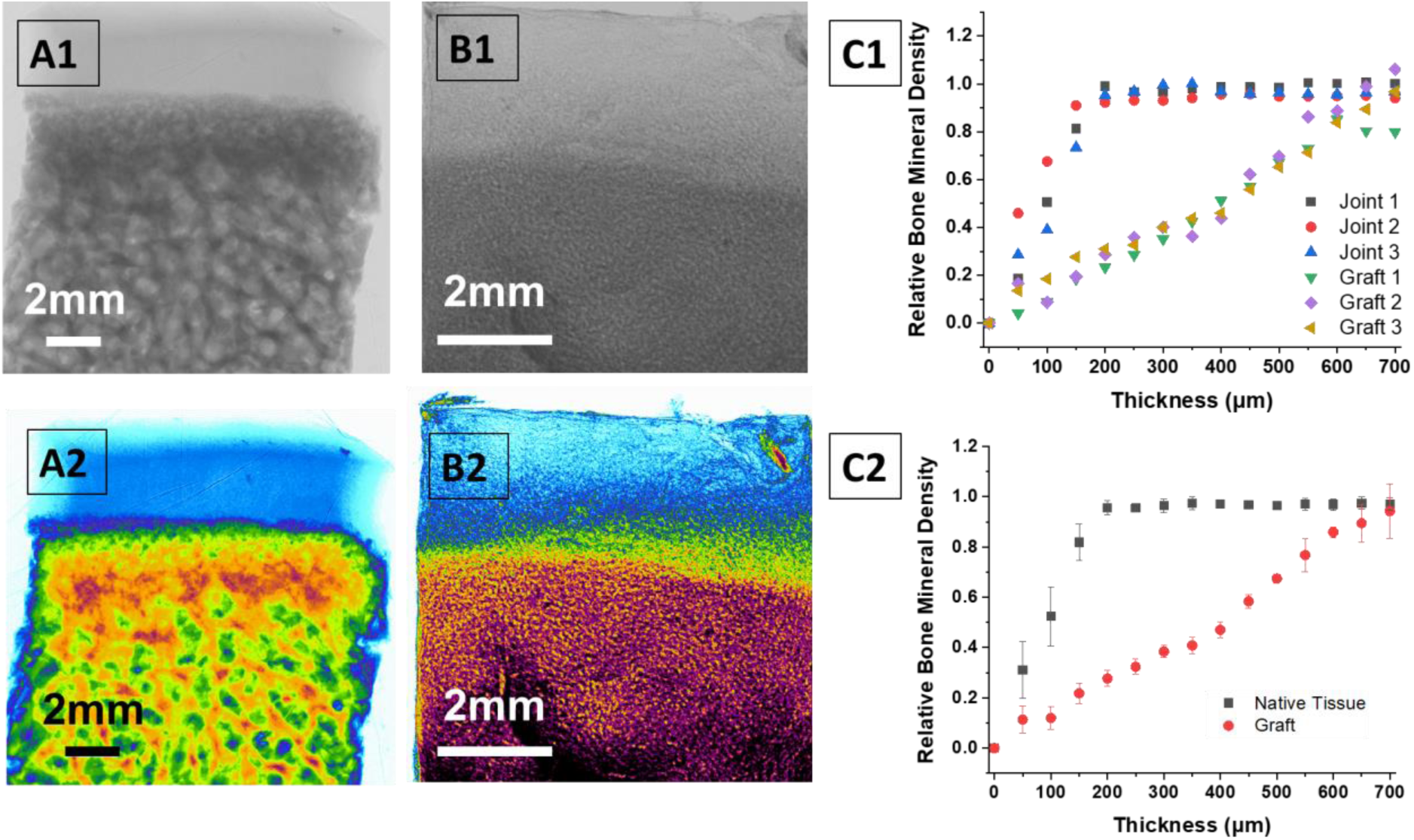
Micro-CT characterization of the OC tissue (A) and the USG (B), and length of grading for the OC tissue (Joint) and the USG (C). The native OC tissue and the graft were visually imaged using micro-CT (A1, B1) and colored for better visualization (A2, B2). The specimens were scanned in layers for the mineral density, processed, and plotted for comparison (C1, C2). Error bars in C2 represent standard deviation.

### 3.5 Bioprinting of hOFBs and chondrocytes

As discussed earlier, a twin screw extruder was used to bioprint the agarose hydrogel containing chondrocytes and agarose/hydroxyapatite hydrogel containing hOFBs. The final cellular USGs were then processed for visualization, with images displayed in Figure 10. As indicated in Figure 10A, the chondrocytes remained viable in Ag hydrogel concentrations ranging from 0.5% to 2%. For the bioprinting process, a 2% Ag concentration was chosen based on its favorable biomechanical properties.

**Figure 10.**
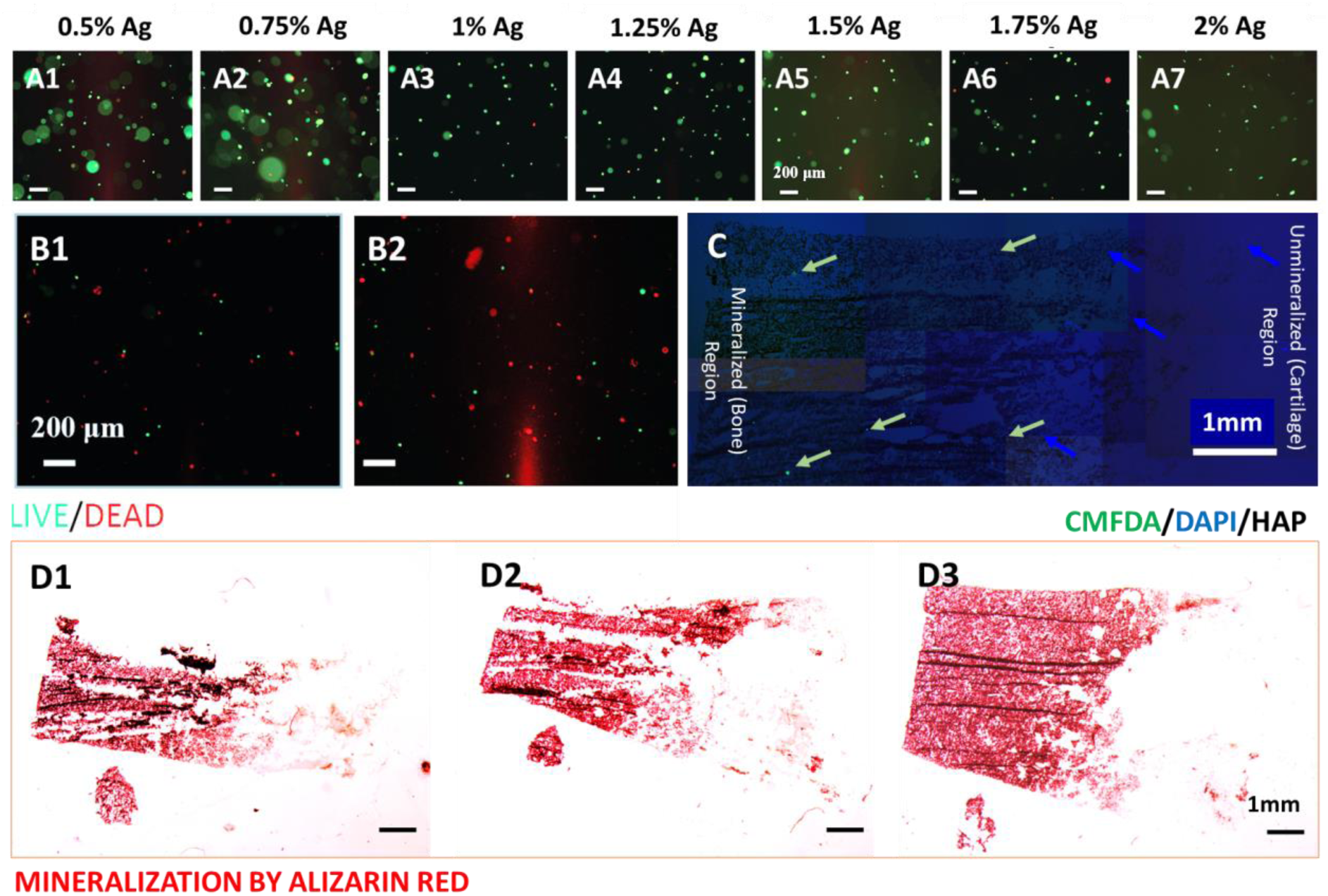
Bioprinting of hFOBs and chondrocytes. (A) Survival (Live/Dead stain) of chondrocytes in Ag gels with different concentrations (0.5-2% w/vol) before bioprinting, (B) Live/Dead staining of chondrocytes (30K cells/mL) in 2% Ag after bioprinting, (C) Stereomicroscope image of CMFDA labeled hFOBs (Green arrows), and (D) Alizarin red stain for mineralized matrix. In C, the green arrows point to CMFDA labeled hFOBs, blue arrows represent DAPI staining, and black color shows HAp. **See the Supplementary Figure at the end for a higher magnification of C**.

Post-bioprinting survival of both chondrocytes and hOFBs was confirmed, as illustrated in Figures 10B and 10C, respectively. Furthermore, the successful graded mineralization within the OC grafts is clearly visible in Figure 10D, demonstrating the effective incorporation and distribution of minerals in the bioprinted structures.

## 4. Discussion

Fabricating scaffolds or grafts that accurately replicate the complex structure and precise mineral gradients of native OC tissue poses significant challenges. The main objective of this study was to engineer a graft that faithfully mimics the mineral gradient observed in native OC tissue. To achieve this, we employed extrusion printing to construct the grafts using agarose solutions with and without hydroxyapatite, which were fed through two separate ports of the extruder. The mineral gradient was carefully engineered by alternately adjusting the flow rates of the agarose solution and the agarose/hydroxyapatite suspension, increasing one while decreasing the other, to ensure a consistent overall flow rate. Several parameters, including the rotational speed of the screws, the die opening, and the timing of flow rate changes, were accordingly adjusted to refine the gradient transition length from 5mm to less than 700 μm. This adjustment allowed us to closely replicate the gradient transition length of the native OC tissue as determined via thermogravimetric analysis. The optimal settings were identified as a rotational speed of 75 rpm, a die opening of 2mm x 2mm, and precise switching between flow rates to achieve the desired gradient.

Thermogravimetric analysis (TGA) was employed to confirm the presence and quantify the content of the hydroxyapatite within the composite USGs. TGA proved especially useful as it can differentiate between organic material decomposition and the residual inorganic content, which primarily consists of hydroxyapatite that remains stable up to 550°C.^15^ The analysis confirmed that approximately 20% of the material did not decompose, indicating the presence of hydroxyapatite.

The linear viscoelastic properties of the USGs were characterized under oscillatory shear and compared with those of native OC tissue. In the tested range, the storage modulus for both the grafts and native OC tissue proved independent of time and temperature. The tan δ values demonstrated that both tissue and the USG exhibited gel-like behavior, with storage moduli significantly exceeding loss moduli, and were frequency-independent. Upon compression, both USGs and native OC tissue showed incomplete stress relaxation over a 200-second period, highlighting the significant role of viscoelasticity in their behavior. Unlike Newtonian fluids, for which the stress decays instantly upon cessation of shearing, or purely elastic materials that maintain constant stress under sustained strain, these materials exhibited typical viscoelastic stress relaxation.

Differences in biomechanical properties between the USGs and native OC tissue were noted. The grafts displayed a consistent, single-slope stress-strain behavior up to failure, whereas native OC tissue exhibited a two-slope stress-strain curve, indicative of two distinct moduli reflecting the cartilage and bone regions.^29^ This suggests a variation in mechanical strength associated with the differences in hydrogel concentration and hydroxyapatite content within the grafts.

Previous efforts to create mineral gradients in osteochondral scaffolds and grafts using unitary structures have faced limitations. For instance, using electrospinning technique, our group (Erisken et al., 2008)^7^ and Liu et al. (2018)^30^ successfully produced thin membranes with mineral gradients of 350 μm and 460 μm, respectively, however, electrospinning produced layered polymer matrices rather than cohesive unitary scaffolds. Other researchers such as Mohan et al.^31^, Singh et al.^32^ and Harley et al.^33^ (from the same research group) developed unitary scaffolds with graded concentrations of minerals and biomolecules but did not specify the gradient transition lengths. Given their use of microparticles ranging from 100-300 μm to form gradients, it is inferred that their gradient transition lengths significantly exceeded 700 μm. Meanwhile, studies by Levingstone et al.^34^, Khanarian et al.^35^, and Erickson et al.^29^ managed to establish gradients with gradient transition lengths exceeding 1mm.

In contrast, our study successfully achieved a gradient transition length of 647±21μm, as determined by TGA, in a unitary graft tailored for OC interface repair and regeneration. This narrow gradient transition length closely mirrors that of the natural OC interface. The bioprinting of chondrocytes and hFOBs into designated zones within the graft demonstrated a physiological resemblance to native OC tissue, affirming cell viability post-bioprinting. This accomplishment underscores the efficacy of the extrusion bioprinting technique in crafting functionally graded grafts suitable for osteochondral tissue engineering. Moreover, the extrusion method offers potential for industrial scale-up and precision manufacturing, enhancing its applicability in clinical settings.

Moving forward, our research will focus on extending the culture period of the bioprinted chondrocytes and hFOBs beyond 30 days to monitor matrix formation and to further analyze the extracellular matrix synthesis by the cells. This continued research will deepen our understanding of the grafts’ behavior in a physiological environment and their potential for clinical applications.

## 5. Conclusion

This investigation aimed to develop unitary synthetic osteochondral grafts that replicate the structural properties of native OC tissue using an innovative extrusion bioprinting technique. This approach was novel in its application of precise bioprinting to create a gradient of mineral concentrations mimicking those found in natural OC interfaces - a key challenge in regenerative tissue engineering.

The primary goal was to fabricate grafts that not only closely match the native OC tissue in terms of mineral gradient transition lengths and viscoelastic properties but also support viable chondrocytes and hFOB cells within their matrix. The use of extrusion bioprinting to achieve narrow gradient transition lengths and integrate living cells directly into the grafts was a novel aspect of this research, pushing the boundaries of current tissue engineering technologies.

One significant challenge was replicating the complex viscoelastic behavior and the precise mineral gradients of natural OC tissue. This was addressed through the controlled extrusion of agarose and hydroxyapatite, allowing for the fine-tuning of material properties and gradient transition lengths. Additionally, ensuring cell viability post-bioprinting was critical and was successfully managed by optimizing the bioprinting parameters to maintain a conducive environment for cell survival and integration.

To advance the clinical application of these bioprinted grafts, several steps are necessary. First, the biomechanical properties of the grafts need enhancement to match or exceed those of native tissues. This could involve experimenting with different material compositions or hybrid structures that incorporate both synthetic and natural materials. Second, long-term in vivo studies are essential to assess the grafts’ performance over time, including their integration with host tissue, the longevity of the repair, and the functional restoration of joint mechanics. Finally, scaling these techniques for clinical trials will require adherence to stringent regulatory standards and effective collaboration between engineers, biologists, and clinicians. This study sets the groundwork for significant potential advancements in orthopedic treatments by providing a method to create grafts that could regenerate complex OC defects and restore joint function, addressing a critical clinical need that affects millions globally.

## Funding Statement

The School of Engineering and Digital Sciences of Nazarbayev University supported the travel expenses for this joint work.

## Conflict of Interest Disclosure

Authors declare no conflict of interest.

---

**Figure 10S:**
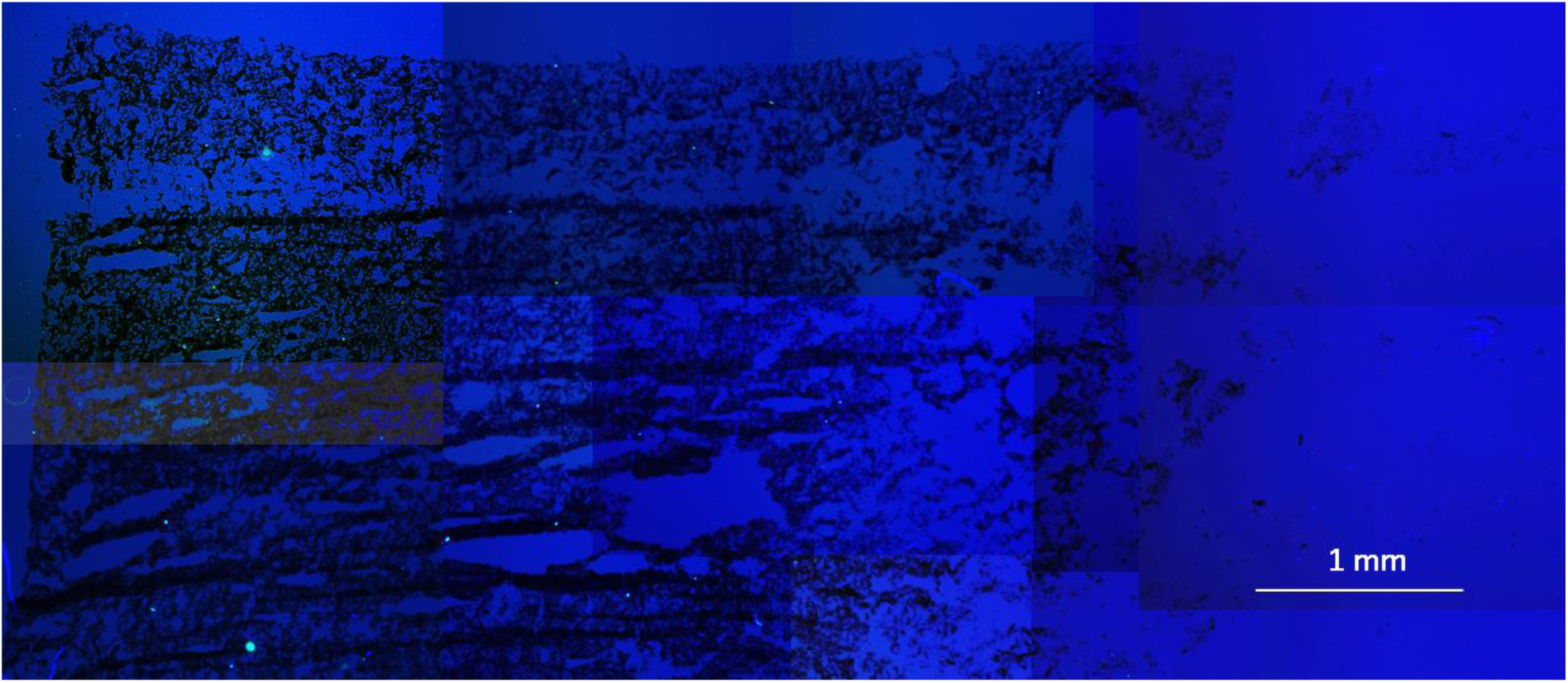
Supplementary Figure

## Notes

### Competing Interest Statement

The authors have declared no competing interest.

